# Effect of flow rate and freezing on cyanocobalamin recovery using a commercial solid phase extraction cartridge

**DOI:** 10.1101/538355

**Authors:** Lauren Lees, Alison M. Bland, Giacomo R. DiTullio, Michael G. Janech, Peter A. Lee

**Affiliations:** College of Charleston, Department of Biology, 66 George Street, Charleston, SC 29424, U.S.A; Hollings Marine Laboratory, 331 Fort Johnson Laboratory, Charleston, SC 29412, U.S.A

**Keywords:** Vitamin B_12_, Multiple Reaction Monitoring, LC/MS, strata-x, flow rate

## Abstract

Analysis of vitamin B12 in sea water is laborious, time consuming, and often requires storage of relatively large-volume water samples. Alleviating these major limitations will increase the throughput of samples and, as a consequence, improve our understanding of the distribution and role of vitamin B12 in the oceans. Previous studies have indicated that target analyte recovery is negatively affected at flow rates exceeding 1 mL min^−1^ using home-made C18 Solid Phase Extraction (SPE) cartridges. In this study, the effect of flow rate on recovery of vitamin B12 was tested across a range of flow rates between 1 and 37 mL min^−1^ using a commercial SPE cartridge containing surface-modified styrene divinylbenzene. Recovery of vitamin B12 at flow rates up to the maximum rate tested did not statistically differ from 1 mL min 1. A second study was conducted to determine whether storage of the SPE cartridges at −20°C had a negative impact on vitamin B12 recovery. Recovery of vitamin B12 from SPE cartridges stored up to 13 days did not differ from unfrozen SPE cartridges. These data suggest that rapid extraction and cold storage of vitamin B12 on commercial SPE cartridges does not negatively affect recovery and offers an economical alternative to field studies.

## Introduction

Cobalamin, commonly referred to as vitamin B_12_, is being increasingly recognized as a limiting nutrient for phytoplankton growth in aquatic environments (Helliwell 2017). Yet, its role in promoting the growth of eukaryotic organisms has long been understood. As early as 1949, growth of the flagellate *Euglena gracilis* was shown to accelerate in the presence of anti-pernicious factor, a.k.a. vitamin B_12_, and was considered as an alternative biological assay for cobalamin deficiency (Hutner et al. 1949).

Numerous forms of cobalamin have been discovered since being first identified in 1948 (Smith and Parker 1948). The majority of the variation in cobalamin forms results from differences in the upper or β-axial ligand on the cobalt ion. While methylcobalamin (MeCbl), (5’-deoxy)adenosylcobalamin (AdoCbl) and hydroxocobalamin (OH-Cbl) are thought to be the main forms of cobalamin in marine microbial communities (Heal et al. 2017; Suffridge et al. 2017), a number of forms, including glutathionyl-, nitroso- and sulfito-cobalamin are known from medical research (Hannibal et al. 2008). Whether or not these additional β-substituted forms are present in the marine environments or microbes remains unknown. More recently, forms of cobalamin produced by bacteria that contain adenine in place of 5,6-dimethylybenzimidazole in the α-axial position (reviewed in (Taga and Walker 2008)) have been identified and called pseudocobalamin. As a result many eukaryotes are unable to acquire pseudocobalamin; thereby, providing yet another point of influence on algal productivity (Heal et al. 2017; Helliwell et al. 2016).

Studies examining the role of cobalamins as a limiting nutrient that controls phytoplankton dynamics in lakes and oceans rely on accurate measurements of cobalamin variants. However, a number of issues exist that severely hamper our ability to assess the impact of cobalamin limitation on phytoplankton growth. First, the concentrations of cobalamins in water samples can approach low femtomoles per liter levels requiring large volumes of water be extracted and/or transported and stored. Transportation of numerous, heavy samples are an obvious strain on logistics and budgets. While recent advances in analytical methodologies have reduced sample volumes from 2-4 L (Okbamichael and Sañudo-Wilhelmy 2004; Panzeca et al. 2009); Panzeca et al. 2009) to 500 mL (Heal et al. 2014; Suffridge et al. 2017), those techniques utilize high-performance liquid chromatography (HPLC) or liquid chromatography-mass spectrometry (LCMS) systems that have relatively large physical sizes and power requirements that limit their use to analytical laboratories far from the field.

Second, recent studies(Heal et al. 2014; Suffridge et al. 2017) have limited the flow of the water sample through SPE cartridges due to the findings of a previously published study(Okbamichael and Sañudo-Wilhelmy 2004) that demonstrated excessive loss of B_12_ at flow rates exceeding 1 mL min^−1^. This latter study used packed columns that were prepared in-house. At a flow rate of 1 mL min^−1^, even a 500 mL water sample would take over 8 hours to extract, which would clearly reduce sample throughput. Third, improving the speed with which samples can be processed reduces the potential for photo-degradation (Juzeniene and Nizauskaite 2013) or the microbial interconversion of the β-axial ligands. In the case of the CarH enzyme, the interconversion is light mediated (Cheng et al. 2016).

With these issues in mind, a series of simple experiments was devised around the use of a commercially available solid-phase extraction cartridge containing a polymer-based sorbent designed to interact with a wide array of chemical properties. The first set of experiments examined the effect of sample flowrate through the SPE cartridge on the retention of cyanocobalamin. In a second set of experiments, cyanocobalamin was extracted from a synthetic seawater matrix and the cartridges stored at −20°C for various lengths of time to assess whether the lightweight and compact SPE cartridges could resist freezing while maintaining retention of the cyanocobalamin. Data suggest higher flow rates and freezing are feasible options for the extraction of cobalamins from field samples.

## Methods

### Chemicals

Mass spectrometry (MS) grade water and acetonitrile (Honeywell) were used for solid phase extraction and LC/MS measurements. All other solvents, acids or bases were of ACS reagent grade or higher.

Stock solutions (1 μmol mL^−1^) of cyanocobalamin (V2876, Sigma) and acetyl-B_12_ (A188028, Toronto Research Chemicals) were made in water and aliquots of 300μl were stored at −80°C. All solutions containing B_12_ were always made or diluted in a five-sided dark box in a dimly lighted laboratory to reduce potential photodegradation. On the day of testing, the vitamin B_12_ stock solution was diluted to a final concentration of 1 μmol mL^−1^ in sterile filtered enriched seawater, artificial water (ESAW) media, pH 8.2.

### Flow Rate Test

Strata-X (2 g/12 mL, Phenomenex) solid phase extraction cartridges containing surface modified styrene divinylbenzene were wrapped in aluminum foil to omit light, conditioned with 10 mL of acetonitrile, and sequentially washed with 10 mL sterile-filtered enriched seawater, artificial water media, pH 8.2 (ESAW;(Harrison et al. 1980)). Vacuum was applied using in-house vacuum (~28 inches Hg gauge) to a vacuum manifold that connected the SPE cartridge by luer-lock connectors. Vitamin B_12_ (1 μmol mL^−1^) was diluted to 1 pmol μL^−1^ in ESAW; and 80μLs was added to 10 mL of ESAW. The ESAW and vitamin B_12_ mixture was vortexed for 5 seconds, and added to the cartridge. Vacuum was applied to the cartridge accordingly to result in the desired flow rate (described below). Following adsorption of vitamin B_12_ with the solid phase, the SPE cartridge was washed with 10 mL MS grade water and allowed to dry under vacuum for 5 minutes. Samples were eluted in 5 mL of 20% acetonitrile in water. The eluate was transferred to a 15 mL amber glass vial and dried using a speedvac on high heat setting for approximately 3-4 hours. The dried samples were stored overnight at −20°C. The next morning, the samples were resuspended at the same time in 400 uL of ammonium formate buffer (0.1% formic acid/0.02% ammonia, adjusted to a pH 6.8 with 25% ammonia (EMD Millipore)) containing acetyl-B_12_ at a concentration of 200 fmol μL^−1^ which was the expected concentration of vitamin B_12_ if recovery was 100%.

Flow rates were first empirically determined prior to the experiment using water with SPE cartridges and opening one, non-connected luer lock valve on the manifold to ambient air. For the experiment, all flow rates were timed using a stop watch and recorded. Any flow rate that did not reside within 15% of the desired flow rate was discarded immediately, which for this test occurred only once for the highest flow rate. Flow rates tested were: (mean ± S.D.) 1.19 ± 0.10, 2.04 ± 0.02, 3.97 ± 0.04, 10.25 ± 0.74, 37.23 ± 3.05 mL min^−^. The highest flow rate determined was the maximum flow rate achieved using the apparatus.

### 1 L Bottle Flow Rate Test

A second analysis was conducted to more closely replicate an actual field sample volume and determine whether recovery of vitamin B_12_ using a high flow rate and larger sample volume differed from the 10 mL volume test samples. A 1 L amber HDPE bottle was filled with 1 L 0.2 μm filtered ESAW containing and 80 μL of vitamin B12 (1 pmol μL^−1^) was added for a final concentration of 80 pmol L^−1^. Two Teflon tubes (0.125 inch OD) were attached to the bottle cap with 0.125 inch nylon bulkhead fittings (Swagelock) that were glued in place using a marine-grade adhesive sealant (3M, 5200). One piece of tubing was fitted through a one-hole, size 0, rubber stopper that was inserted into the top of the conditioned SPE cartridge. The second tube was taped to the side to act as a snorkel when the bottle was inverted and relieve pressure in the bottle as sample was being vacuumed through the SPE cartridge. To achieve high flow rates a 1 L filtering flask with a rubber stopper containing a single luer-lock port was used. The flask was connected to a Savant VP100 two-stage pump (displacement of 3.5 CFM, ~20 mTorr maximum vacuum) for filtration using vacuum tubing. The 1 L bottle was inverted and suspended above the flask using a ring stand. Maximum vacuum was applied to achieve the highest flowrate possible with the apparatus (30.26 ± 4.04 mL min^−1^). The extraction was replicated four times. Samples were dried and resuspended as described above.

### Effect of freezing and storage time on Vitamin B_12_ Recovery

Solid phase extraction, vitamin B_12_ concentration and loading, and reagents were identical to those described above unless otherwise noted. Vitamin B_12_ in 10 mL ESAW media were adsorbed to SPE cartridges at a flow rate of approximately 30 mL min^−1^ using the a vacuum manifold with in-house vacuum (~28 inches Hg gauge). Twenty-four SPE cartridges were sequentially adsorbed with the vitamin B_12_ in ESAW. Four cartridges underwent elution to provide a time-zero data point against which frozen SPE cartridges would be compared. The remaining cartridges were stored in a laboratory freezer at −20°C. During the SPE filtration, start times were noted on a wall clock in the laboratory. Each group of four replicates was adsorbed within 20 minutes of the first cartridge within the respective group. Freezing times were 1 day (24 hours), 2 days (48 hours), 7 days (168 hours), and 13 days (312 hours). On the day that the SPE cartridges were removed from the freezer, the time was noted on the same wall clock and recorded. Processing start times were within 15 minutes of noted duration. Upon removal from the freezer, SPE cartridges were washed with 10 mL MS grade water and dried under vacuum for 5 minutes. Vitamin B_12_ was eluted with 5 mL of 20% acetonitrile. The samples were dried and resuspended exactly as described above for the flow rate test.

### Multiple Reaction Monitoring (MRM)

All data were acquired using a Xevo-TQS triple quadrupole mass spectrometer (Waters) coupled to an M-Class UPLC (Waters). Samples (100 μL) were placed into amber, glass autosampler vials and loaded into the autosampler set at 10°C. A volume of 1 μL sample was injected into a 5μL loop. Vitamin B_12_ sample was loaded at 12 μL min^−1^ directly onto a C18 column (Acquity UPLC M-Class T3 Spherical Silica, 300 μm × 150 mm, 1.8 μm) using Buffer A (0.1% formic acid/0.02% ammonia in water, pH 3.5) and held at Buffer A for 2 minutes to allow the sample to equilibrate. Buffer B (acetonitrile) was introduced into the linear gradient from 0-15% over 4 minutes and held at 15% for 2 minutes before increasing to 80% for 2 minutes. The column was re-equilibrated for 4 minutes using 100% Buffer A. The total run time was 17 minutes. Column temperature was set to 35°C. The sample was ionized in a Waters ZSpray™ ESI source using a low flow probe with the following source parameters: source temperature, 150°C; desolvation temperature, 200°C; capillary voltage, 3.0 kV; cone voltage, 121V; cone gas flow, 150L Hr^−1^; desolvation gas flow, 800L Hr^−1^; collision gas flow, 0.15 mL Hr^−1^; nebulizer gas flow, 7bar. Data were collected in positive ion mode. The following monitored transitions with collision energies in parentheses from the +2 charged masses were generated using the Intellistart feature of MassLynx: vitamin B_12_ 678.57 -> 147.17 (40v), 678.57 -> 359.21 (24v), 678.57 -> 456.91 (34v); acetyl vitamin B_12_ 699.60 -> 147.17 (40v), 699.60 -> 401.18 (24v), 699.60 -> 686.04 (18v). Scanning was conducted at unit resolution and dwell times were set to 0.019s. Standard curves were generated to ensure linearity of the response between 25 fmol to 500 fmol on column. Data were exported to Targetlynx for regression and quantification. Because standard concentration slopes were linear and were not significantly different from each other (slope T-test, P=0.99) Vitamin B_12_ concentrations were estimated by: [(peak area of vitamin B12/ peak area of acetyl vitamin B12) × concentration of acetyl vitamin B12.

### Statistical Analysis

Normality and equal variance of data was tested using the Shapiro-Wilk test and Equal Variance Test features built into SigmaPlot v.11, respectively. Multiple group comparisons were tested using one-way ANOVA and post-hoc comparisons using the Holm-Sidak test. Groups were considered different when p<0.05.

## Results

All experiments were designed to provide measurements (200 fmol μL on column, S/N >5,000) well above the lower limit of quantification (0.995 fmol μL^−1^ on column), estimated using embedded calculations of peak-to-peak noise within TargetLynx. Coefficient of variation from triplicate injections within batches was less than 1%. The highest carry-over measured, based on injection of buffer A alone between sample injections, was less than 0.03%.

Individual recovery of vitamin B_12_ measured across flow rates from approximately 1 to 37 mL min^−1^ ranged between 72% and 112%. Mean (± S.D.) values for the different flow rate categories were: 1ml min^−1^, 108 ± 6%; 2ml min^−1^, 82 ± 7%; 4ml min^−1^, 88 ± 5%; 10ml min^−1^, 97 ± 4%; 37ml min^−1^, 102 ± 3% (Figure 1). Although the highest recoveries were noted at the lowest flow rate 1 mL min^−1^), this difference was not significant (ANOVA, Holm-Sidak test, p=0.43) compared to the highest flow rate (37 mL min^−1^). Coefficient of variation across all flow rates were less than 10% (Figure 1), but were notably lower at the highest flow rates compared to the lowest flow rates. Interestingly, recovery at the 2 mL min^−1^ flow rate was significantly lower than all but the 4 mL min^−1^ group, which could not be explained by obvious deviations in protocol or instrument performance. Based on this comparison, flow rates up to 37 mL min^−1^ perform equal to 1 mL min^−1^ based on recovery using a commercially prepared SPE cartridge with polymeric resin. An attractive advantage of higher flow rates is that sample preparation is considerably hastened.

**Figure 1.**
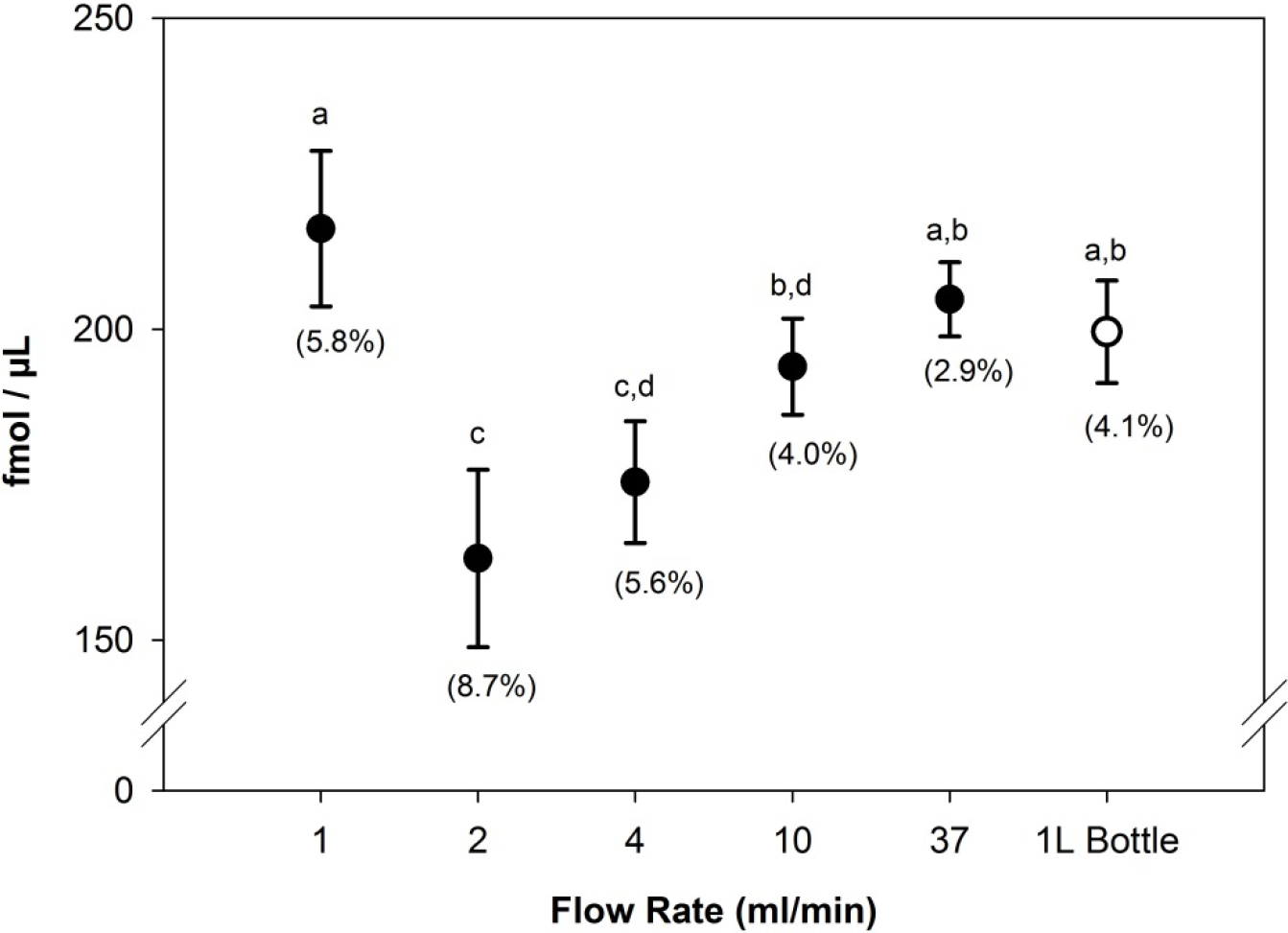
The effect of flow rate across the SPE cartridge on measurements of vitamin B12 (cyanocobalamin). Data are reported as mean ± standard deviation. Vitamin B12 measurements reported are for a 1 μL injection on the analytical C18 column. The expected measurement is 200 fmol μL^−1^. Coefficient of variation is given for each flow rate group in parentheses. Closed circles denote groups that belong to the small volume extraction test set using 10 mL ESAW media. The open circle denotes extraction of vitamin B_12_ from 1L of ESAW media using high vacuum. N=4 per group. Differences between groups are indicated by letter code where groups containing identical letters are not different and *vice versa* (One-way ANOVA, Holm-Sidak, p<0.05).

To determine whether high recovery and low variability could be maintained using a sample volume that more closely replicated those collected from field sites, vitamin B_12_ was measured from 1 L spiked-ESAW media. Because high flow rates could not be achieved using a simple manifold with house vacuum, a dedicated flask and more powerful vacuum pump was included for this test. With this design, the total amount of time needed to filter 1 L of ESAW was approximately 33 minutes, which considerably shorter compared to the estimated time it would take at a slower flow rate of 1 mL min^−1^ (~16.7 hours). Recovery of vitamin B_12_ from this experiment ranged between 94 to 103% with a mean recovery of 100 ± 4.1%. These data were not significantly different compared to the 1, 10 and 37 mL min-^1^ flow rates in the small volume comparison (**Figure 1**), suggesting SPE adsorption at high flow rates is applicable to larger volumes and is durable with regard to high vacuum pressures.

### Cold Storage

Compared to the unfrozen time zero cartridges, freezing and thawing did not appear to negatively affect the commercial SPE cartridge. No cracking of the plastic housing or loss of solid phase during washing and elution was observed. Individual recoveries from this experiment ranged from 90 to 119% and no differences were detected between any groups (Figure 2). Coefficient of variation ranged from 5.3 to 10.5% and relative variability did not trend with time. These data suggest that extraction of seawater vitamin B_12_ on to SPE cartridges with subsequent freezing for storage and thawing for analysis is a practical alternative to storing large volumes of sea water over short periods of time. Further experiments are underway to assess the viability of freezing and storing the cartridges over longer periods of time.

**Figure 2.**
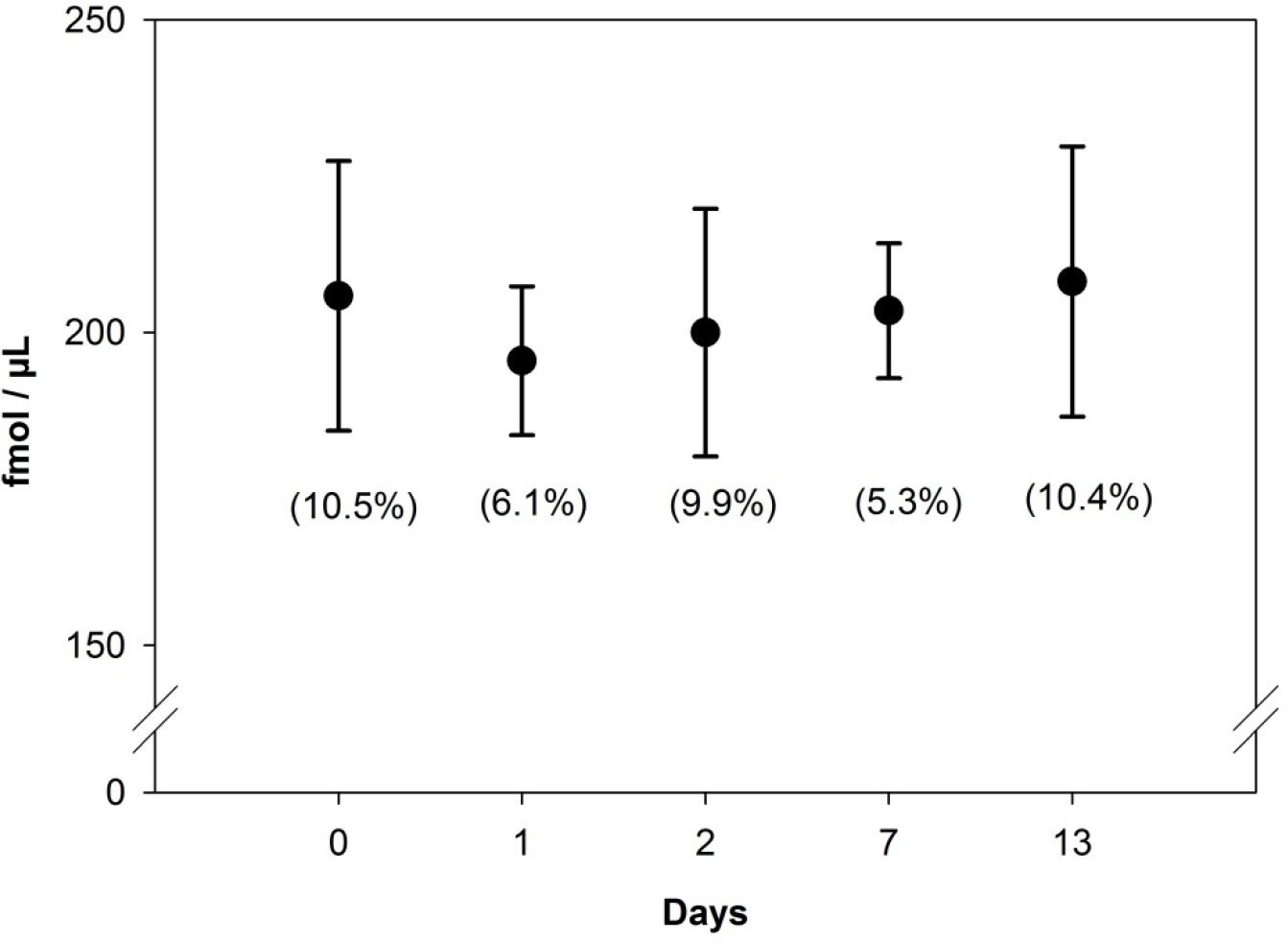
The effect of cold storage and time on measurements of vitamin B12. Data are reported as mean ± standard deviation. Vitamin B_12_ measurements reported are for a 1 μL injection on the analytical C18 column. The expected measurement is 200 fmol μL^−1^. Coefficient of variation is given for each flow rate group in parentheses. N=4 per group. There were no statistical differences between groups (One-way ANOVA, Holm-Sidak, p<0.05).

## Discussion

With interest into the influence of cobalamin on phytoplankton growth and resource competition growing, there is a greater need to rapidly measure cobalamin or pseudocobalamin species at low concentrations. Solid phase concentrating devices are a popular choice due to their strong retention of target analytes and ability to selectively elute target compounds to remove potential matrix suppressors. In the ocean, cobalamins are often found at sub-picomolar levels(Heal et al. 2014; Suffridge et al. 2017) making these difficult to quantify in small sample volumes; thereby, leading to the need for collecting and extracting large sample volumes or injecting larger volumes of SPE processed samples onto the C18 column. Based on current protocols, the flow rate often cited for C18 SPE extraction of cobalamin is 1mL min^−1^ (Okbamichael and Sañudo-Wilhelmy 2004). In this study, we used a similar, but not identical solid phase and found that 1mL min^−1^ provides excellent recovery and there is no issue with laboratory protocols that have published with this low flow rate. However, the data provided in this study suggest that the time required for extracting cobalamins can be greatly hastened up to flow rates of 37 mL min^−1^ without a loss in recovery. For a 1L water sample, the extraction can be done in 1/30^th^ the time necessary, thus allowing multiple, larger volume samples to be extracted in excess of 1L within a daily routine.

To address the issue of sample collection at remote field stations or on oceanographic expeditiions where electricity and cold storage are available, but where high-end mass spectrometers are unlikely to be located and storage space is limited; we tested whether SPE columns could be frozen and whether an adsorbed analyte could be recovered. Freezing of the SPE cartridges did not appear to impact the recovery vitamin B_12_ as there were no differences detected over time. Variability was not different compared to the unfrozen cartridges. Although SPE cartridges were only tested for 13 days, the recovery of vitamin B_12_ was within acceptable limits and there was no indication that further time at cold storage would have impacted the recovery, but the effect of long-term storage should be verified if samples will be stored longer than 2 weeks.

Because this study was not a comprehensive test comparing multiple commercial SPE phases, nor did the study utilize actual field samples, the conclusions reached here should serve as a rationale to test flow rate and freeze storage conditions using investigator-preferred SPE cartridges, be it made in the laboratory or from a commercial source. The benefits are several: 1) overall reduction in processing time; 2) significant reduction in shipping weight; 3) significant reduction in storage space; 4) reduction in sample injection volume.

## Acknowledgments

This work was funded in part through the U.S. National Science Foundation (OCE-1428915, OCE-1436458 and AOE-1644073). Any opinions, findings, and conclusions or recommendations expressed in this material are those of the author(s) and do not necessarily reflect the views of the National Science Foundation. This article is contribution number TBD to the Graduate Program in Marine Biology at the University of Charleston, Charleston, South Carolina.

## Author contribution

All authors contributed to the writing of the manuscript. LL constructed the filtration apparatus, conducted the SPE experiments, and assisted in data analysis. AB conducted the mass spectrometry analysis and collated data for analysis. JD consulted on the experimental design and analysis. MJ conceived of the study design and oversaw the experiments in the laboratory. PL co-planned the experiments, constructed the 1L bottle, and analyzed the data.

## Literature Cited

Cheng, Z., H. Yamamoto, and C. E. Bauer. 2016. Cobalamin’s (Vitamin B(12)) Surprising Function as a Photoreceptor. Trends in biochemical sciences 41: 647–650.

Hannibal, L., A. Axhemi, V. Glushchenko Alla, S. Moreira Edward, E. Brasch Nicola, and W. Jacobsen Donald. 2008. Accurate assessment and identification of naturally occurring cellular cobalamins, p. 1739. Clin Chem Lab Med.

Harrison, P. J., R. E. Waters, and F. J. R. Taylor. 1980. A broad spectrum artificial sea water medium for coastal and open ocean phytoplankton. J Phycol 16: 28–35.

Heal, K. R. and others 2014. Determination of four forms of vitamin B12 and other B vitamins in seawater by liquid chromatography/tandem mass spectrometry. Rapid Commun Mass Spectrom 28: 2398–2404.

Heal, K. R. and others 2017. Two distinct pools of B12 analogs reveal community interdependencies in the ocean. Proc Natl Acad Sci U S A 114: 364–369.

Helliwell, K. E. 2017. The roles of B vitamins in phytoplankton nutrition: new perspectives and prospects. The New phytologist 216: 62–68.

Helliwell, K. E. and others 2016. Cyanobacteria and Eukaryotic Algae Use Different Chemical Variants of Vitamin B12. Curr Biol 26: 999–1008.

Hutner, S. H. and others 1949. Assay of Anti-Pernicious Anemia Factor with Euglena. Proceedings of the Society for Experimental Biology and Medicine 70: 118–120.

Juzeniene, A., and Z. Nizauskaite. 2013. Photodegradation of cobalamins in aqueous solutions and in human blood. Journal of photochemistry and photobiology. B, Biology 122: 7–14.

Okbamichael, M., and S. A. Sañudo-Wilhelmy. 2004. A new method for the determination of Vitamin B12 in seawater. Anal Chim Acta 517: 33–38.

Panzeca, C. and others 2009. Distributions of dissolved vitamin B12 and Co in coastal and open-ocean environments. Estuarine, Coastal and Shelf Science 85: 223–230.

Smith, E. L., and L. F. Parker. 1948. Purification of anti-pernicious anaemia factor. Biochem J 43: viii.

Suffridge, C., L. Cutter, and S. A. Sañudo-Wilhelmy. 2017. A New Analytical Method for Direct Measurement of Particulate and Dissolved B-vitamins and Their Congeners in Seawater. Frontiers in Marine Science 4.

Taga, M. E., and G. C. Walker. 2008. Pseudo-B12 joins the cofactor family. J Bacteriol 190: 1157–1159.

